# Postglacial migration across a large dispersal barrier outpaces regional expansion from glacial refugia: evidence from two conifers in the Pacific Northwest

**DOI:** 10.1101/2020.01.05.895193

**Authors:** MC Fernandez, FS Hu, DG Gavin, G deLafontaine, KD Heath

## Abstract

Understanding how climate refugia and migration over great distances have facilitated species survival during periods of past climate change is crucial for evaluating contemporary threats to biodiversity. In addition to tracking a changing climate, extant species must face complex, anthropogenically fragmented landscapes. The dominant conifer species in the mesic temperate forests of the Pacific Northwest are split by the arid rain-shadow of the Cascade Range into coastal and interior distributions, with continued debate over the origins of the interior populations. If the Last Glacial Maximum extirpated populations in the interior then postglacial migration across the arid divide would have been necessary to create the current distribution, whereas interior refugial persistence could have locally repopulated the disjunction. These alternative scenarios have significant implications for the postglacial development of the Pacific Northwest mesic forests and the impact of dispersal barriers during periods of climate change. Here we use genotyping-by-sequencing (ddRADseq) and phylogeographical modeling to show that the postglacial expansion of both mountain hemlock and western redcedar consisted largely of long-distance spread inland in the direction of dominant winds, with limited expansion from an interior redcedar refugium. Our results for these two key mesic conifers, along with fossil pollen data, address the longstanding question on the development of the Pacific Northwest mesic forests and contrast with many recent studies emphasizing the role of cryptic refugia in colonizing modern species ranges.

**Statement of Significance:** Understanding whether habitat fragmentation hinders range shifts as species track a changing climate presents a pressing challenge for biologists. Species with disjunct distributions provide a natural laboratory for studying the effects of fragmentation during past periods of climate change. We find that dispersal across a 50-200-km inhospitable barrier characterized the expansion of two conifer species since the last ice age. The importance of migration, and minimal contribution of more local glacial refugia, contrasts with many recent studies emphasizing the role of microrefugia in populating modern species distributions. Our results address a longstanding question on the development of the disjunct mesic conifer forests of the Pacific Northwest and offer new insights into the spatiotemporal patterns of refugial populations and postglacial vegetation development previously unresolved despite decades of paleoecological studies.

## Introduction

In response to the environmental changes during periods of rapid climate change, populations must migrate to remain within their niche, tolerate the changes through adaptation, or go extinct (1–3). However, ongoing anthropogenic climate change has the potential to outpace both migration and adaptation, threatening a loss of genetic diversity in rear-edge populations or even extinction (4–7). The current distributions of many plant species have been shaped by past periods of climate change, particularly during the Quaternary ice ages (8, 9). For some species, northern glacial refugia facilitated *in situ* persistence and seeded the rapid postglacial colonization of newly suitable habitat (10–12), while postglacial migration over great distances allowed other species with distant refugia to expand across the landscape (13). Thus, studying how species have migrated in response to past climate change can yield insights into their long-term capacity for migration under anthropogenic warming and help provide more robust predictions of their future (14).

In contrast with past periods of climate change, however, current range shifts must also contend with complex, human-modified landscapes, in which the risks of climate change are compounded by habitat loss and fragmentation due to widespread agricultural and urban development (15, 16). Fragmented landscapes reduce dispersal, isolating existing populations and hindering the invasion of newly favorable habitats (17), even for wind dispersed plant species (18). Nevertheless, models of future species responses to climate change tend to neglect human land-use effects and assume homogenous landscapes (19, 20).

Species with disjunct populations provide a naturally fragmented distribution that can act as an approximation for the effects of anthropogenic land-use. A modern species disjunction is the result of either recent (e.g., postglacial) long-distance migration or the long-term persistence of remnants from a bygone species range. The former underscores the ability of populations to migrate across complex landscapes in response to climate change, whereas the latter implies that small refugial populations may be critical for species perseverance under shifting climatic conditions. Past demographic events leave telltale genetic imprints on extant populations, allowing inference of the colonization sources and migration routes of intraspecific lineages (8, 21). It is therefore possible to use patterns of genetic variation to distinguish between refugial persistence and recent migration (21). Genetic methods, consequently, overcome the low capacity of paleorecords to detect small populations (22).

The mesic temperate forests in the Pacific Northwest (PNW) of North America provide an ideal biogeographical model to explore the consequences of complex landscapes on Quaternary climate-induced migration. The ranges of the dominant conifer species, and dozens of other mesic-adapted species, in these forests are split by the arid rain-shadow of the Cascade Range into coastal and interior distributions. Since the pioneering work of Daubenmire (23), the postglacial origin of the modern interior distribution has been studied for several taxa (24–26), but remains a subject of debate for the dominant conifer species. If the climatic conditions of the Last Glacial Maximum (LGM; ∼21 ka BP) extirpated populations in the interior, necessitating postglacial migration across the rain shadow divide from coastal populations to form the modern interior range, we would anticipate genetically similar distributions. Alternatively, if local persistence in the interior developed into a modern range independent of the coast, we expect genetically distinct interior and coastal distributions as a result of genetic drift during their prolonged glacial isolation from one another. These alternative scenarios have significant implications for the postglacial development of PNW conifer forests and whether fragmented habitat presented a surmountable barrier to the expansion of its species. Here we present phylogeographic insights from next-generation sequencing studies of two mesic conifers with disjunct distributions to address the relative importance of ice age refugial persistence versus long-distance postglacial migration in creating the modern range, and therefore infer their ability to migrate over complex landscapes.

## Results and discussion

We gathered genome-wide marker data for two wind dispersed and late-successional mesic conifer species common to the disjunct forests of the PNW: mountain hemlock (*Tsuga mertensiana*; > 95% outcrossing) (27) and western redcedar (*Thuja plicata*; 71.5% outcrossing) (28). We used genotyping-by-sequencing (ddRADseq) (29) to resolve SNP markers for 154 western redcedar individuals (22 populations) and 149 mountain hemlock individuals (21 populations) (Table S1). A combination of Bayesian clustering analysis (*Structure*)(30), principal component analysis (PCA), discriminant analysis of principal components (DAPC), and approximate Bayesian computation modeling (*DIYABC*)(31) allowed us to infer the population history of these species.

Bayesian clustering (Fig. 1), PCA (Fig. 2), and DAPC (Fig. S1) each support two distinct genetic clusters in mountain hemlock and four in western redcedar. Mountain hemlock is divided into northern and southern clusters, which meet in southern Washington state with little indication of admixture (Fig. 1a,b). Western redcedar is comprised of a large central cluster extending across Washington and southern British Columbia, with substantial admixture between this central cluster and three smaller clusters in the regions of Idaho, northern California, and the Haida Gwaii islands (Figure 1c,d).

**Figure 1.**
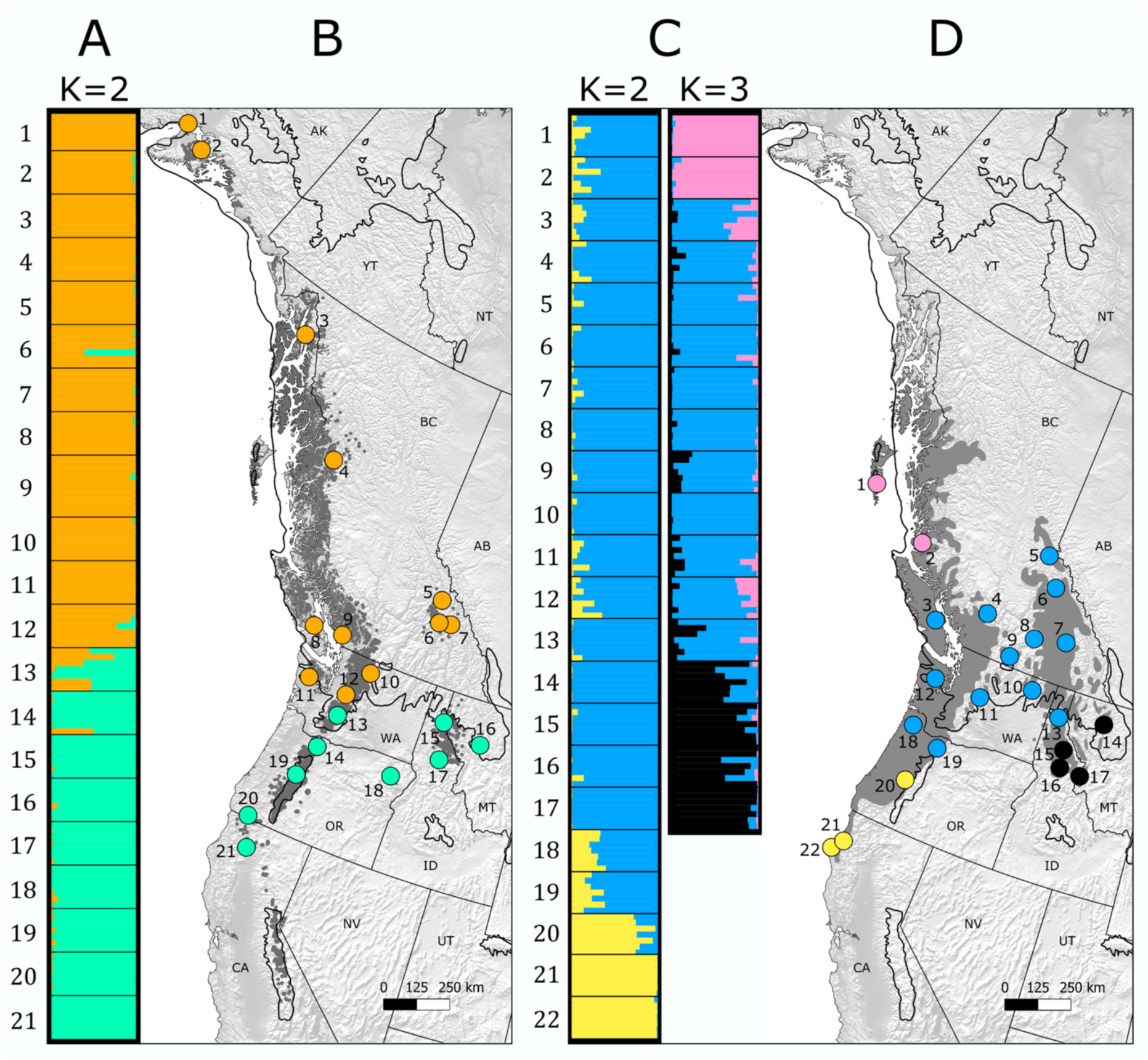
Genetic structure indicative of glacial refugia. The modern distributions of the two study species (left, mountain hemlock; right, western redcedar) are indicated with a darker shade of grey and the maximum extent of the ice age glaciers is indicated with a black line. (A) Structure results of 21 mountain hemlock populations (K=2: orange = north cluster; teal = south cluster). (B) Spatial patterns of hemlock Structure results. (C) Hierarchical Structure results of 22 western redcedar populations with full dataset (K=2; yellow = south cluster, blue = north cluster) and without the southern cluster populations (K=3; black = Idaho cluster, blue = Central cluster, pink = Haida Gwaii cluster). (D) Spatial patterns of redcedar Structure results (combined, K=4).

**Figure 2.**
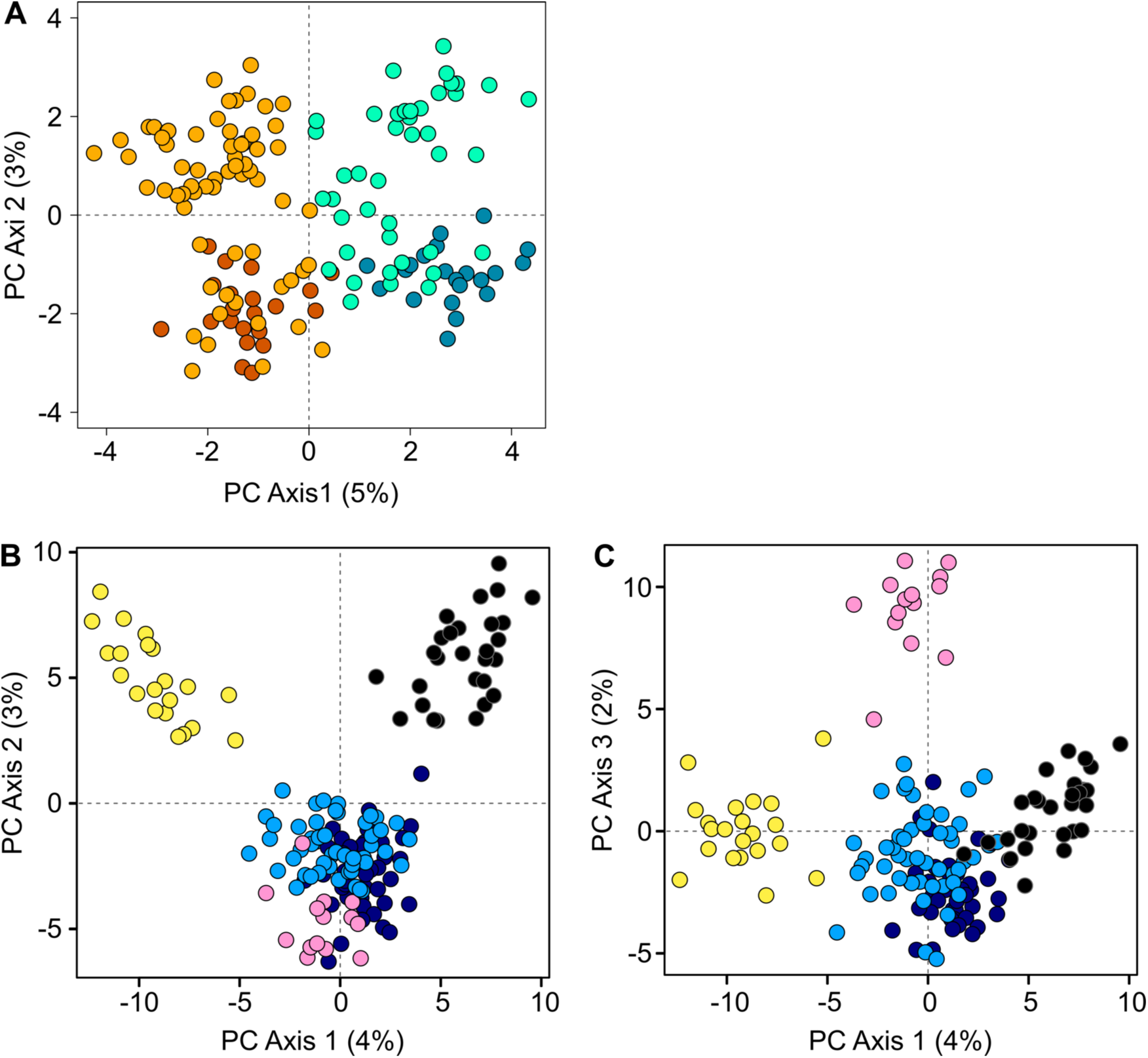
Genetic differentiation of inferred refugial groups. PCA results for mountain hemlock (**A**) and western redcedar (**B**,**C**). Data points are colored according to their *Structure* genetic cluster association. (**A**) Mountain hemlock principal component analysis (K=2; orange/dark orange = south cluster, blue-green/teal= north cluster). The coastal and interior distributions of each lineage is indicated with the lighter and darker colors, respectively. (**B**) The first and second axes of the western redcedar principal component analysis (K=4; yellow = California cluster, black = Idaho cluster, light blue/dark blue = Central cluster, pink = Haida Gwaii cluster). The coastal and interior distributions of the Central cluster are indicated with the lighter and darker colors, respectively. (**C**) The first and third axes of the western redcedar principal component analysis, with the same color indicators as in 2b.

For mountain hemlock, the north/south genetic division in the interior distribution mirrors that of the coastal populations (Fig. 1a,b). This pattern, in conjunction with the lack of appreciable genetic divergence between the coast and interior (Fig. 2a), suggests that each of the two lineages (north/south) independently migrated into the interior along separate dispersal routes. Indeed, an analysis of molecular variance (AMOVA) indicated that significant genetic variation exists between the northern and southern lineages as opposed to non-significant genetic variation between the coast and interior distributions (Table S2). Although there is some indication of partial segregation between coastal and interior sites along the second PCA axis (Fig. 2a), a subset of coastal sites and the interior sites of each lineage overlap substantially (Fig. 1b; North: sites 10-12, South: sites 13-14). At *K*=4, DAPC supports the same grouping (Fig. S1c). Geographically, the coastal sites that cluster with the interior are adjacent to previously-hypothesized mesic migration corridors between the coastal and interior distributions (26), suggesting that these coastal locations were sources of hemlock migrants into the interior. While habitable patches may have been available in an otherwise inhospitably arid rain shadow to facilitate their dispersal inland, exhaustive mapping has not identified such patches in the modern landscape (32). Despite recent dispersal into the interior, there is incomplete infilling of mountain hemlock into its potential distribution between the separately-established northern and southern interior disjunctions; this may have been a result of limited dispersal both across patchy mountain-tops in a direction orthogonal to the dominant winds, and from upwind coastal areas (32).

Paleorecords and paleoclimate simulations suggest that the modern mesic climate only emerged in the interior between 6.6 – 4 ka (33–35). Patterns of mountain hemlock presence in the pollen record indicate exclusively south-coastal populations at the LGM and emergence in the interior only within the last 5,000 years (Fig. S2) (32), consistent with genetic inferences of postglacial inland migration. Likewise, the best (Fig. 3a) of six ABC models (Fig. S3a) of the demographic history of hemlock indicates a significantly older median age of division between the north and south lineages (ca. 99 ka, 95% CI = 77–135 ka; Table S3) relative to the comparatively recent coastal/interior divisions within each lineage (North: ca. 11 ka, 95% CI = 6–14 ka; South: ca. 25 ka, 95% CI = 9–46 ka; Table S3), with bottlenecks in both interior populations. The modeled age of the north-south divergence suggests that the divergence occurred prior to the LGM followed by ice age population isolation, whereas the more recent divergence between coastal and interior populations likely supports postglacial migration.

**Figure 3.**
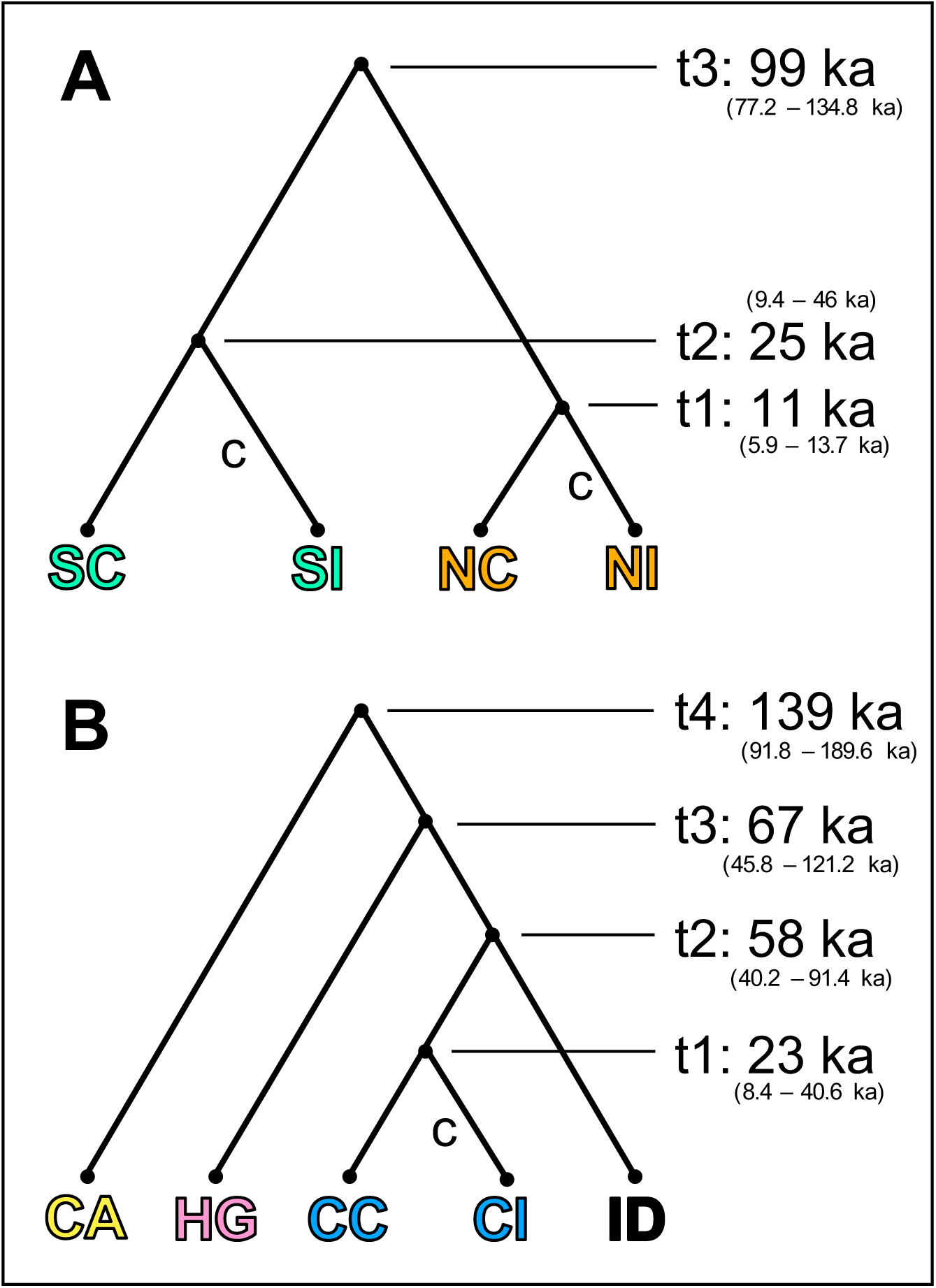
Inferred phylogenetic tree structure and divergence dates. The divergence dates presented are median rounded values. **(3a**) Best mountain hemlock *DIYABC* model (NC = coastal north cluster, NI = interior north cluster, SC = coastal south cluster, SI = interior south cluster, “**c**” = reduced population size). (**3b**) Best western redcedar *DIYABC* model (ID = Idaho cluster, CC = coastal Central cluster, CI = interior Central cluster, HG = Haida Gwaii cluster, CA = California cluster, “**c**” = reduced population size).

For western redcedar, a similar lack of genetic divergence between the coastal and interior distributions suggests that most of the current interior populations are coastal in origin. However, in contrast with mountain hemlock, there is evidence of a redcedar refugial lineage unique to the interior (Fig. 1c,d). As in hemlock, an AMOVA indicates that grouping sites by their genetic lineages accounts for more genetic variation than grouping by coast and interior distribution (Table S2). Nevertheless, there was a small, but significant, amount of variation partitioned between the coast and interior for western redcedar. The first two axes of the PCA for redcedar indicates separation into three distinct groups (Fig. 2b), a southern cluster in California, an interior cluster in Idaho, and a central cluster of the populations found between the other two. The central cluster further separates the northernmost coastal populations along the third principal component (Fig. 2c). Much like the hemlock lineages the “Central” redcedar lineage extends across both the coastal and interior populations with no significant genetic distinctiveness between the two distributions (Fig. 1d; 2b,c). Separation into *K*=3-5 DAPC clusters shows the same patterns of sampling-site grouping as the PCA (Fig. S1). *Structure* results corroborate these patterns of population structure at corresponding *K* values (Fig. 1c).

Modeling the demographic history of redcedar using DIYABC supports the inferred species history. Our best (Fig. 3b) of six ABC models (Fig. S3b) estimates the oldest median age of divergence among lineages separating the California cluster from the three more northern clusters (ca. 139 ka). The three lineages composing the northern clade diverged from one another more recently around the same time during the last glaciation (ca. 58-67 ka). Haida Gwaii is estimated to have diverged first (ca. 67 ka), followed closely by the Idaho and “Central” clusters (ca. 58 ka). The interior populations of the “Central” cluster diverged from coastal populations comparatively late (ca. 23 ka) and experienced a bottleneck. These relative age estimates suggest a north/south divergence during the last interglacial period, glacial isolation among the northern lineages, and postglacial dispersal from the coastal “Central” populations into the interior.

Unlike mountain hemlock, the morphologically indistinct nature of redcedar pollen grains (Cupressaceae family) and very infrequent macrofossil evidence limit the paleoecological data available with which to corroborate our genetic inferences. Abundant evidence suggests a cold-maritime climate for western Oregon and Washington during the LGM (36, 37) and hindcasted species distribution models suggest redcedar was restricted to coastal SW Oregon and NW Oregon (38). Additionally, a long pollen record from the unglaciated portion of the interior distribution in Idaho showed a rapid rise in abundance of Cupressaceae pollen after ca. 6.3 ka following its complete absence for the previous >100 ka, suggesting only a recent regional establishment of western redcedar regardless of origin (39).

Our genetic clusters are generally congruent with a previous microsatellite study of redcedar (40); however, our extensive geographic sampling together with genome-wide SNPs and demographic modeling allowed us to identify a previously-unknown interior refugium and infer divergence times. Evidence of interior refugial persistence in redcedar correspond with several amphibian species (41) and several forest herbaceous species associated with redcedar understories (42), though other species show indications of recent dispersal inland (26). The presence of an interior western redcedar refugium suggests that habitats with mesic conditions must have persisted in Idaho during the last ice age, and by extension such areas may have harbored many endemic or disjunct mesic-adapted species (43).

Several factors might explain why, despite the presence of western redcedar populations in an interior refugium in the heterogeneous terrain of central Idaho, coastal migrants played the predominant postglacial role in founding the current interior distribution of the species. First is timing: the widespread establishment of a mesic climate on the coast likely preceded that of equivalent latitudes in the interior by thousands of years (33–35), providing coastal redcedar populations a substantial head start in their expansion. Moreover strong westerlies in the PNW postglacial air mass circulation likely facilitated the inland expansion of wind dispersed conifers (44). Second is location: it is unlikely that the Idaho redcedar refugium overlapped with much of the current species distribution, as Herring & Gavin (39) inferred tree-less periglacial conditions at the glacial maximum from a site near the current distribution’s southern edge. Instead, the refugium may have persisted further south and/or been confined within canyons harboring small mesic habitats, constraining postglacial expansion northward (25, 42, 45). Indeed a higher concentration of understory and endemic species in the southern portion of the modern interior distribution supports southern-interior mesic refugial persistence as well as slow, incomplete postglacial dispersal northwards by those mesic species (26). Finally, expansion from a refugium may also involve complex interactions and competition between species attempting to pioneer the newly available mesic habitat, critically slowing incipient expansion from the cryptic interior refugium relative to that of the heavy propagule pressure from the much larger coastal distribution (46).

Surprisingly, our study suggests that the fragmented landscapes of the Pacific Northwest did not present major limitations to the postglacial migration and range expansion of these two important mesic conifers. Between the two species, there is evidence that three lineages independently crossed a substantial migration barrier, with one redcedar lineage even outcompeting the local interior population. Recent studies have emphasized the importance of cryptic refugia as source populations for postglacial expansion (10), as they could help explain the rapid rates of plant migration inferred from the paleorecord (47, 48). Although refugial populations have undoubtedly had a significant impact for some species, our redcedar data suggest that the importance of migration cannot be downplayed even in the presence of local refugia. Although fossil data are less vulnerable to false positives (11), our results also underscore the importance of genetic methods when interpreting the significance of any particular refugium in postglacial expansion dynamics. In our case, detecting the presence of an interior western redcedar refugium in the paleorecord would not have meant that it was the source population for forest expansion within the interior, or that long-distance migration from the coast was unimportant.

We infer remarkable migration capacities for our study species, potentially indicating substantial long-distance dispersal events if we assume that the disjunctions were established by wind dispersal and that the width of the modern minimum dispersal barrier (ca. 200 km and 50 km for hemlock and redcedar, respectively) was not reduced during different climatic periods following deglaciation. The assumption that the dry rainshadow persisted over the Holocene is supported by the glacial geomorphology in Idaho, which indicates a predominant westerly flow of moist airmasses, as today, during the late-glacial period (49). Furthermore, a trend of increasing available moisture through the Holocene, which is evident throughout the Pacific Northwest (e.g.,35), is not consistent with greater connectivity between the coast and interior in the early-than the late-Holocene. Given this, how are such dispersal distances possible? Previous studies have shown that the vast majority of wind dispersed conifer seeds fall near the parent tree, within several hundred meters for mountain hemlock (50, 51) and approximately one hundred meters for western redcedar (52–54). Vertical updrafts greatly increase dispersal distances and simulations suggest that > 1% of seeds with a similar terminal velocity may be uplifted above the canopy into turbulent winds (55). Although dispersal distances > 1 km are very difficult to study using mechanistic simulation models or with direct experimental evidence (56), support for such long-distance dispersal is provided by other conifer species with disjunct populations. For example, on oceanic islands in the Pacific Ocean at least four intra-specific disjunctions > 100 km of anemophilous conifer species exist for which single dispersal events must be invoked (57). Comparable dispersal distances have been inferred for disjunct populations of ponderosa pine (58), though birds are known dispersers of large-seeded pines while wind is the only known disperser of our study species.

In contrast to the Idaho refugium, the redcedar refugium on the north coast of British Columbia may have played an important role in postglacial range-filling. Substantial evidence exists that portions of coastal British Columbia were ice-free during the last glaciation and served as a refugium for numerous species (59). Although there is no paleoecological data supporting the persistence of trees on the islands of Haida Gwaii, habitat modeling has suggested a suitable glacial climate for both study species (24). Our results imply a potential Haida Gwaii refugium for western redcedar, and that the population succeeded in expanding onto the mainland during the postglacial, where it would have encountered migrants northbound from the “Central” refugial lineage (60). Unlike the migration into the interior disjunction, however, a western redcedar refugium in Haida Gwaii would not necessarily have encountered substantial migration barriers. Low ice age sea level exposed the Hecate Strait during the Pleistocene, providing contiguous habitable terrain connecting Haida Gwaii with the mainland (59) that could have supported redcedar migration during deglaciation. Therefore, Haida Gwaii migrants would not have faced dispersal challenges as substantial as those of coastal migrants into the interior distribution.

Although we suggest that our study species adeptly navigated complex landscapes in the past, facilitated by their ability to migrate (61), climate change still threatens populations of these and other more vulnerable species. The same broad habitat tolerance that enabled redcedar to persist in tenuous northern glacial refugia in the past (62) leads to model-based predictions of future stability for redcedar, but precipitous decline for hemlock given its vulnerability to the anticipated loss of subalpine habitat (63, 64). In other words, even incredible migration capabilities become irrelevant when no suitable habitat remains (65). Our work highlights how species’ tolerance of variable climates and migration ability across fragmented landscapes ultimately interact to determine how their ranges shift through time.

## Methods

Samples were collected from 21 populations of mountain hemlock and 22 populations of western redcedar (Table S1). At each sampling site, foliage from each of 6-10 trees was collected (pairwise distance between individuals >100 m), preserved in silica desiccant, and stored at 4°C at the University of Illinois at Urbana-Champaign. DNA was extracted from 20 mg of unblemished foliage using a modified CTAB procedure (66) for hemlock and the Qiagen DNeasy Plant Mini Kit for redcedar. Floragenex Inc. (Portland, Oregon USA) prepared ddRADseq libraries (67) using *PstI*/*MseI* enzymes and sequenced twelve lanes of single-end Illumina HiSeq 2000/2500, for a total of 728.1 million reads (filtered to 655.9 million) for 149 mountain hemlock samples and 846.9 million reads (filtered to 678.2 million) for 154 western redcedar samples. We processed raw sequencing reads using the program *Stacks* (v1.35) and identified 30,007 variable hemlock loci and 40,701 variable redcedar loci present in at least two populations. *Stacks* demultiplexed the reads, filtered for quality and barcode/RADtag presence, and categorized the data into the most statistically probable SNPs and genotypes (68, 69).

To decipher genetic population structure, we used the Bayesian clustering program *Structure* (v2.3.4; 30) with the *StrAuto* utility program (v0.3; 70). We used the *Stacks* sub-program *Populations* to output a *Structure*-formatted dataset of putatively unlinked SNPs by selecting a single SNP from each locus. For a hemlock locus to be included in this analysis, we required it to be present in at least 7 of 21 sampling sites and in ≥75% of the individuals at each of those sites, which resulted in 1,158 loci. Redcedar loci required presence in 20 of 22 sampling sites and ≥75% of the individuals, resulting in 872 loci. We ran *Structure* using the admixture model, with correlated allele frequencies and no prior population information. The analysis consisted of 20,000 burn-in steps and 100,000 replicates of 1-5 genotypic groups (*K*), each of which was run 10 times. The output was compiled using *Structure Harvester* (71) and optimal *K* evaluated based on LnP(K) and Δ*K* (72) and visual inspection of each *K* for indications of geographically correlated clustering. To evaluate indications of additional population structure in our western redcedar *Structure* run, we conducted an additional *Structure* run of only the northern cluster. The hierarchical run required presence in 15 of 17 northern sampling sites and ≥75% of the individuals, resulting in 1661 loci for analysis.

We additionally visualized the population structure of our genetic data using principal component analysis (PCA) (73) and discriminant analysis of principal components (DAPC) (74). These methods do not rely on the assumptions underlying *Structure* (e.g., Hardy-Weinberg equilibrium and linkage disequilibrium) (74) and provide an alternative evaluation of population structure. These analyses were run using the *adegenet* (v2.0.1) (75) package in *R* (v3.2.0) (76). We ran a PCA to infer axes of genetic variation in our data and visualized them based on our *Structure* results and patterns of geographically-correlated clustering. We used DAPC to infer the number of populations by partitioning genetic variance ‘between-group’ and ‘within-group’ and identifying the configuration that maximizes discrimination between the two (74). To run these two analyses (PCA, DAPC) we utilized a high-quality dataset of 310 hemlock loci (at least 50% of a population and 20 of 21 populations) and 364 redcedar loci (at least 75% of a population and 21 of 22 populations), with a single SNP per locus. Any missing data was imputed with the most common genotype at that site.

We conducted an analysis of molecular variance (AMOVA) (77) to estimate the molecular variance among our sampling sites and evaluate competing demographic scenarios (e.g., ancient vicariance and recent colonization). Because this analysis is sensitive to missing data, we generated a reduced dataset containing only 14 populations in hemlock and 17 populations in redcedar (interior sites and coastal sites at comparable latitudes) using the *Stacks* subprogram *Populations* to maximize the number of loci common to all populations. For a locus to be included in this dataset, we required it to be present in all sampling sites and in ≥50% of the individuals at each site, producing 292 hemlock loci and 12,019 redcedar loci, with a single SNP per locus. We ran the AMOVA in Arlequin (v3.5) (78) with 10,100 permutations to examine the distribution of genome-wide genetic variation between groups (*F*_CT_), among populations within groups (*F*_SC_), and among populations (*F*_ST_).

We used the approximate Bayesian modeling program *DIYABC* (v2.0) (31) to evaluate potential demographic scenarios and estimate divergence times between the genetic clusters in our study species. Genetic clusters spanning both the coastal and interior ranges were divided into separate groups to infer evidence of postglacial dispersal. We utilized high-quality datasets of 209 hemlock loci (≥65% of a group, 4 of 4 groups) and 183 redcedar loci (≥90% of a group, 5 of 5 groups), with a single SNP per locus. Due to the complex interactions of the four western redcedar genetic clusters and their admixture, only genetically distinct redcedar populations with ≥85% assignment to a single *Structure* cluster were included in this analysis (98 individuals from 17 sites). Please see the supplemental for additional methods.

Pollen-based vegetation reconstructions from the PNW provide spatial and temporal context for interpreting phylogeographic data. We conducted a review of published paleorecords that include mountain hemlock pollen to trace its patterns of glacial persistence and postglacial migration based upon the presence/absence of hemlock pollen through time. Mountain hemlock pollen can be readily identified at the species level (79) and as a consequence is a common component of Pacific Northwest sediment core pollen descriptions. We obtained published pollen records from the Neotoma Paleoecology Database (http://www.neotomadb.org/), as well as additional paleorecords where available. Mountain hemlock is well known for its tendency to leave weak pollen evidence in the fossil record, even on occasions where it was prevalent in the local watershed (80–83). We therefore considered hemlock to be locally established when its pollen percentage reached at least 1% of the counted total, indicating its presence within 140 km (32). Records were then classified into five time slices (>20 kyr, >15 kyr, >10 kyr, >5 kyr, and >0 kyr) according to the basal age of the record and the age at which mountain hemlock first reached the 1% pollen threshold. The timeframe of mountain hemlock pollen presence was based upon the author’s evaluation of the paleorecord’s reliable age range. To account for chronological uncertainty, we included a record in an age category if pollen evidence of mountain hemlock occurred within 200 years of the time specified (e.g., the ‘>20 kyr’ age class includes records 19,800 years and older). Because many pollen records have >1% hemlock pollen at the oldest (i.e., basal) sample, this survey may not necessarily capture the earliest arrival of the species at all sites and should thus be interpreted as the minimum age of local hemlock establishment.

## Acknowledgements

We would like to thank Urban M, Lauren L, Chang W, Blyth G, Weintraub L, Napier J, Chipman M, Fernandez JM, Fernandez CL, Fernandez VI, Blitvic N, Burke P, Russell J, Ferguson C, and the Cowichan Lake Research Station for their field and laboratory assistance. Funding was provided by NSF (DEB-1146017, and NSF-GRFP to Fernandez M), and the Graduate College at the University of Illinois.

