## Supplemental Figures and Tables for "Postglacial migration across a large dispersal barrier outpaces regional expansion from glacial refugia: evidence from two conifers in the Pacific Northwest"

### Supplementary Information for

*Title*: A tale of two conifers: Evidence of postglacial long-distance migration of western redcedar and mountain hemlock across fragmented landscapes

*Author Affiliation*: Fernandez MC[1], Hu FS[1], Gavin DG[2], deLafontaine G[3], and Heath KD[1]

[1]University of Illinois at Urbana-Champaign; Department of Plant Biology
[2]University of Oregon; Department of Geography
[3]Université du Québec à Rimouski; Département de biologie, chimie et géographie

**This PDF file includes:**

Figs. S1 to S3
Tables S1 to S4

**Supplemental Methods, DIYABC**:

In order to incorporate the phylogeny of the western redcedar genetic clusters inferred by *Structure*, PCA, and DAPC, we generated a phylogenetic tree with which to inform our *DIYABC* model. We used the *Stacks* sub-program *Populations* to output a genepop-formatted dataset with sites grouped according to the majority genetic clusters assigned. The dataset was a single SNP from each locus and required presence in 4 of 4 redcedar groups and ≥80% of individuals per group, resulting in 4,120 loci. We used the program *TreeFit* (1) to assemble a population tree based on the pairwise genetic distances (2) between populations using a neighbor-joining method. Support for the tree was evaluated based on 1,000 bootstrap permutations, all of which reproduced the same tree. The tree provided an *a priori* assumption from which to base our modeling of demographic scenarios and divergence times in *DIYABC*. Based on the more prominent north-versus-south division between California and the other lineages in our $K$=2 *Structure* results (Fig. 1c), we chose to root the tree with the California lineage for our demographic modeling in *DIYABC*.

Six demographic scenarios for population expansion and contraction were tested for each species with broad parameter settings (Fig. S3). The preferred scenario was then re-analyzed with parameter settings constrained to 95% of data values inferred by the model in the first stage of analysis. For each scenario, one million datasets were simulated and their summary statistics compared to those observed to evaluate their likelihood. We selected scenarios based upon the program's logistic regression and 'direct approach' methods of comparison, as well as the goodness-of-fit model checking function (3). The median posterior distribution estimates of divergence times were used to annotate the phylogenetic tree (Table S3). Although western redcedar trees are able to begin producing seeds in as little as 10 years of age under well-lit conditions (4, 5), production normally begins at around 20-30 years of age (6). Mountain hemlock trees mature in a similar manner, generally beginning to bear seeds at around 20 years of age (7). Because divergence times are calculated in terms of generations, we assumed a conservative 20-year generation time to estimate approximate divergence date for both species.

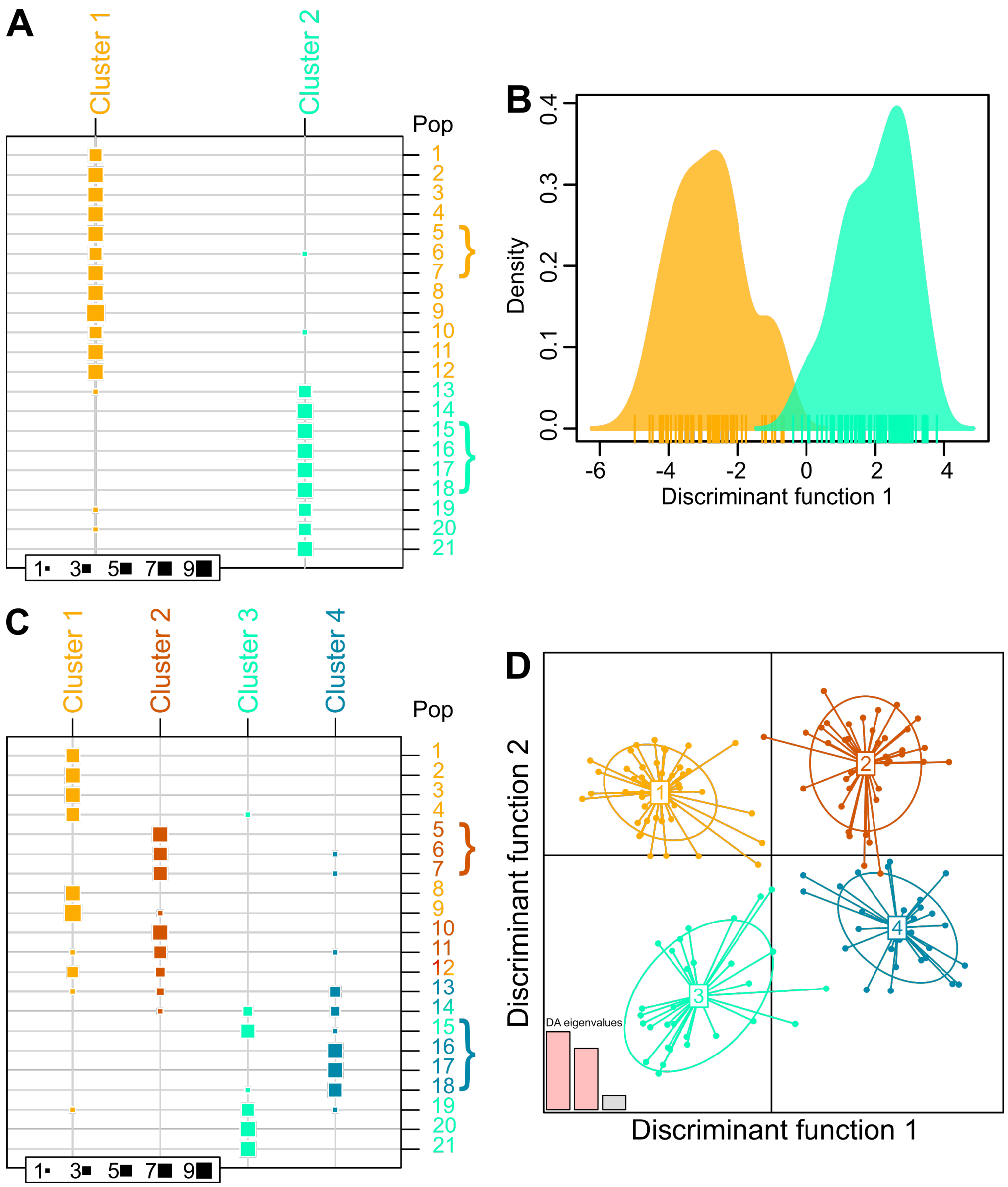


**Fig. S1.** Discriminant analysis of principal components (DAPC) of genetic variation. (See below).

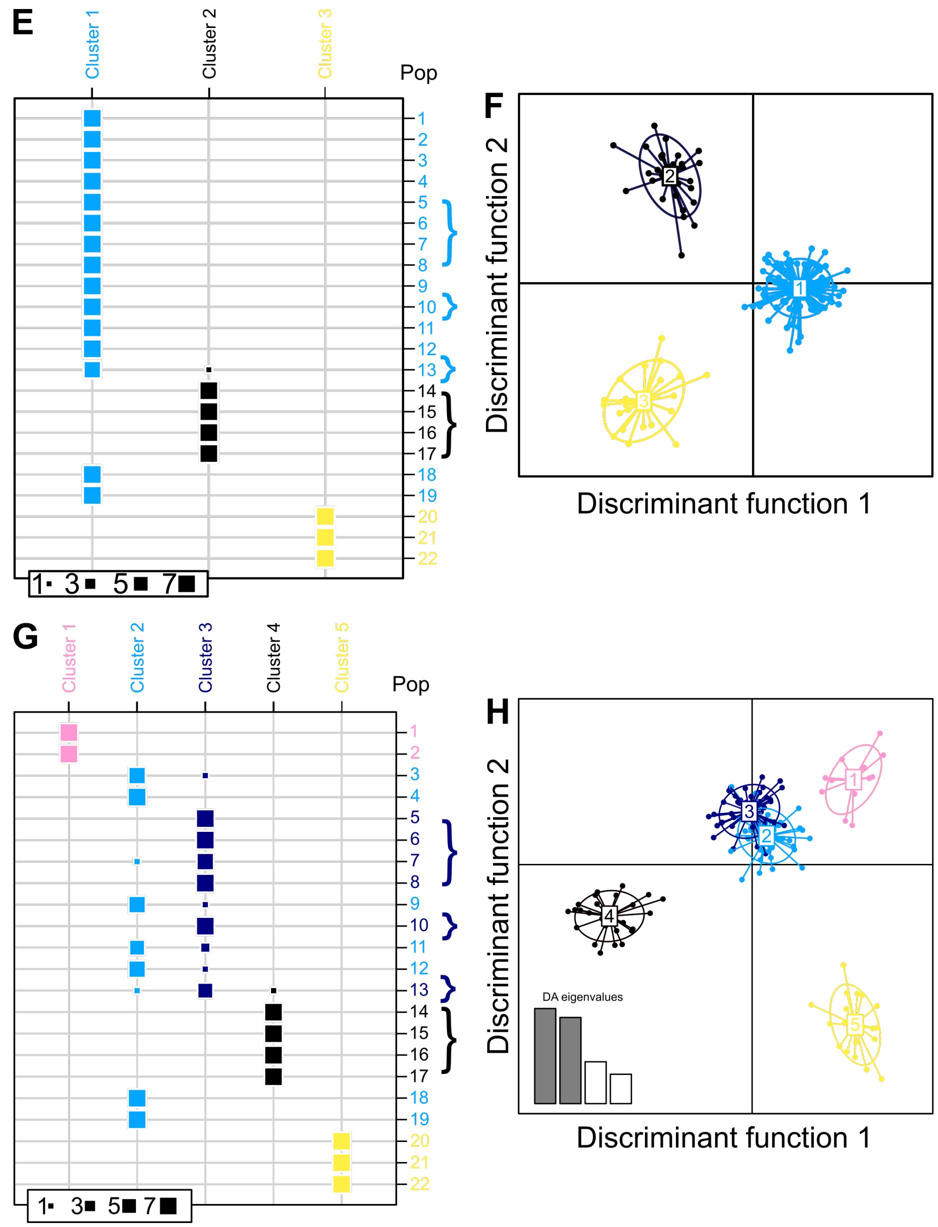


**Fig. S1. (Continued)** Discriminant analysis of principal components (DAPC) of genetic variation. Based on the number of clusters (*K*) requested, the analysis attempts to separate the genetic data into the most differentiated groups possible. The listed population numbers correspond to those in Figure 1a,c and square-size indicates the number of individuals (1-7) from a given population that are categorized as a member of each cluster. Interior distribution populations are marked with brackets. We demonstrate hemlock genetic differentiation into *K*=2 (**A**) and *K*=4 (**C**) clusters as seen in our PCA results. Any further *K* above this value lose geographical coherence as distinct groups. The separation of both analyses discriminant functions are also shown (**B**,**D**). We also demonstrate redcedar genetic differentiation into *K*=3 (**E**) and *K*=5 (**G**) clusters as seen in our PCA results. Weak segregation and substantial overlap between the coastal and interior populations of the central cluster are visible in *K*=5 (clusters 2 and 3). The separation of both analyses discriminant functions are also shown (**F**,**H**).

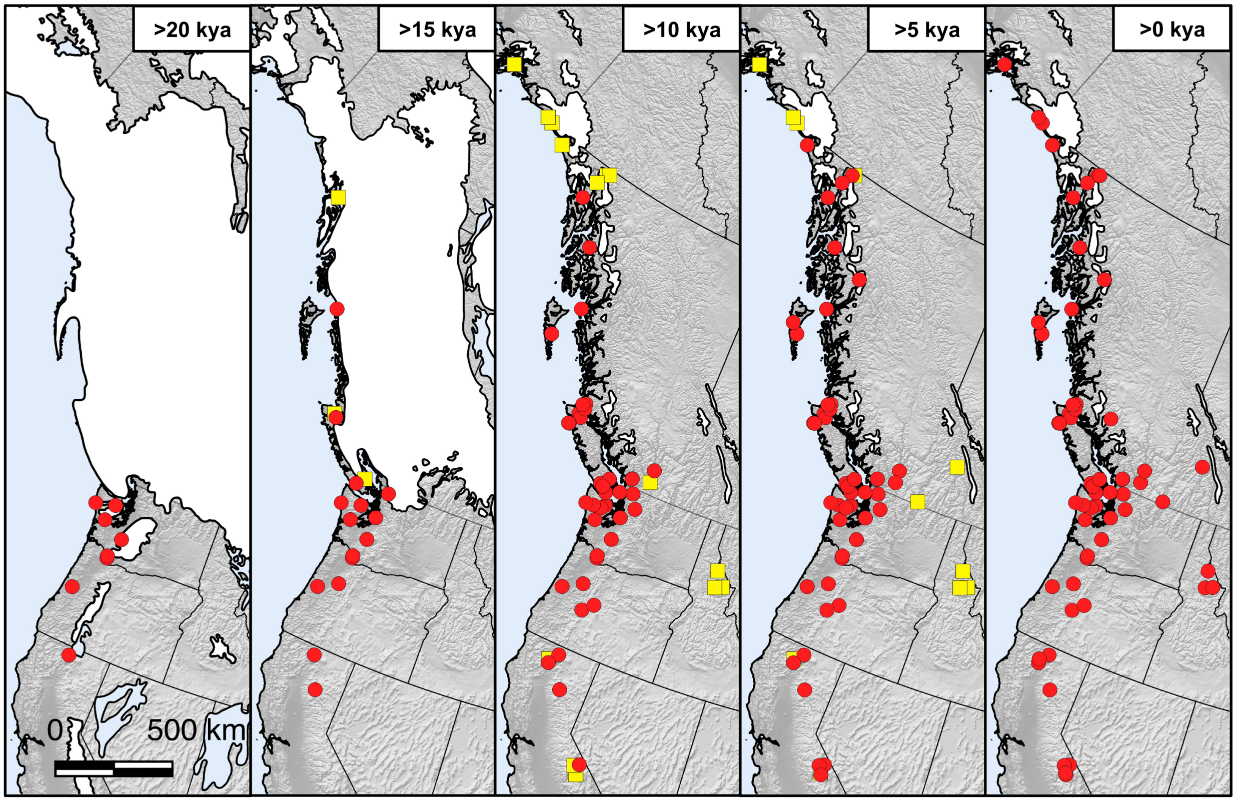


**Fig. S2.** Survey of radiocarbon-dated published pollen records that record presence of mountain hemlock pollen. The pollen records have been partitioned into five time steps, separated by 5,000 year intervals, to provide a visual representation of the emergence of mountain hemlock across the Pacific Northwest according to the paleorecord. A yellow square indicates that a pollen record includes data for this time step, but that mountain hemlock pollen had not yet reached the 1% representation threshold in this, or a previous, time step. A red circle indicates that during this, or a previous, time step mountain hemlock reached at least 1% representation in the pollen record. Pollen records indicate the presence of mountain hemlock along much of the west coast of the United States during the last glacial maximum (>20 kya time step), including more northern sites near the ice sheets in Washington state. In the next four time steps (>15 kya to >0 kya), mountain hemlock emerges in a south-to-north pattern up into Alaska as the ice sheets retract. Pollen records within the interior distribution (northern Idaho and eastern British Columbia) that record mountain hemlock pollen are all 10,000-15,000 years in age and have no indication of mountain hemlock pollen until the final time step (>0 kya). One site in northern Idaho (not shown but located 12-km south of the modern range) had no mountain hemlock pollen recorded for more than 120,000 years (Herring and Gavin, 2015).

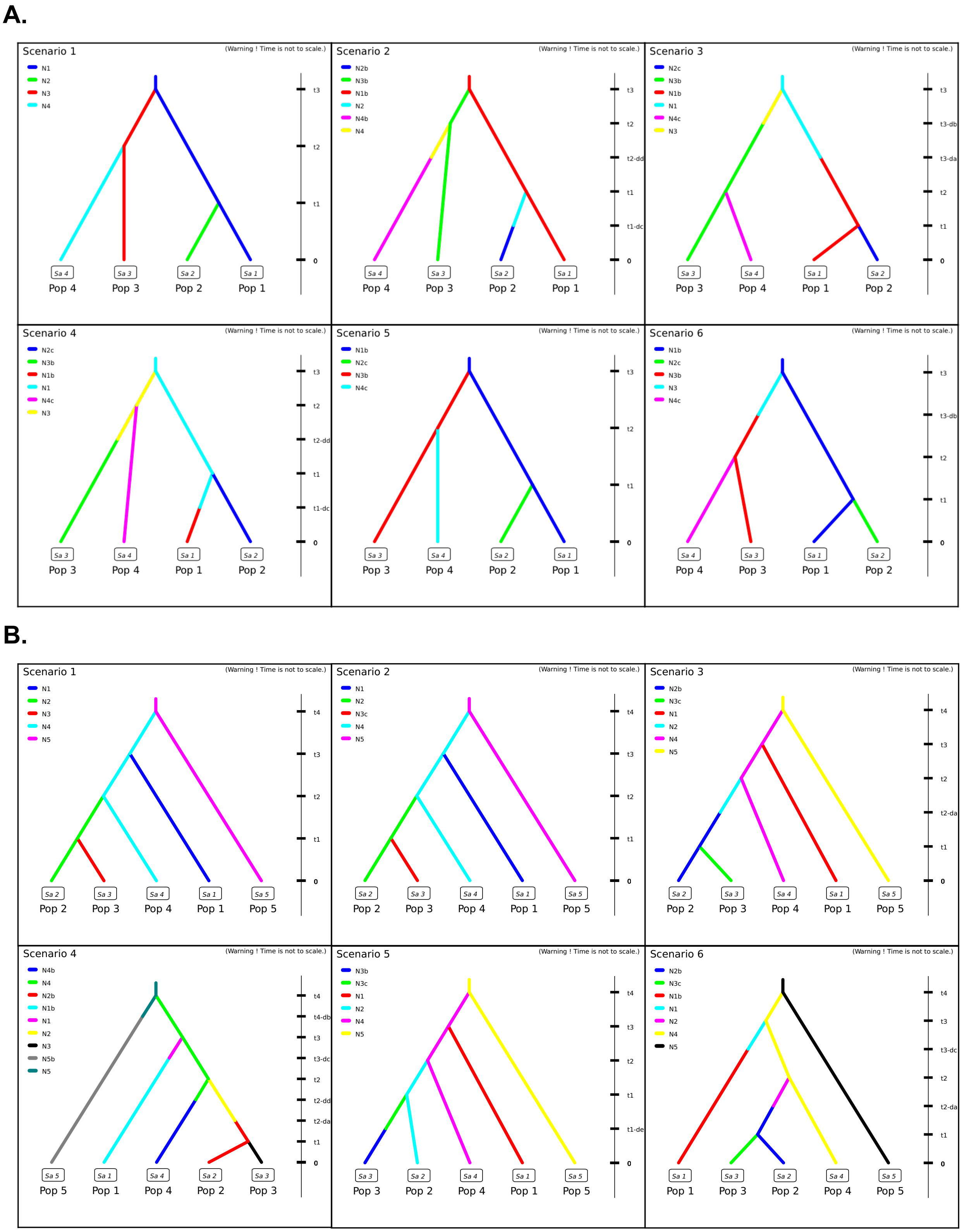


**Fig. S3.** Hemlock demographic scenario for recent dispersal inland (See below).

**Fig. S3.** Hemlock and Redcedar demographic scenarios for dispersal inland (Continued).

Demographic scenarios for dispersal inland. DIYABC results for the mountain hemlock and western redcedar datasets, performed following to the program's standard operating manual. (**A**) Demographic scenarios tested for mountain hemlock. N1 and N2 indicate the coastal and interior populations, respectively, of the southern genetic cluster. N3 and N4 indicate the coastal and interior populations, respectively, of the northern genetic cluster. An appended "b" indicates a set parameter of increased effective population size compared to the base population, whereas an appended "c" indicates a set parameter of decreased effective population size compared to the base population. (**B**) Demographic scenarios tested for western redcedar. N1 indicates the Haida Gwaii genetic cluster, N2 and N3 the coastal and interior populations, respectively, of the "central" cluster, N4 the Idaho genetic cluster, and N5 the California genetic cluster. An appended "b" indicates a set parameter of increased effective population size compared to the base population, whereas an appended "c" indicates a set parameter of decreased effective population size compared to the base population.

**Table S1.** Sample collection information.
Table of populations for each species and a GPS coordinate representative of the region foliage samples were collected from (see methods for sampling strategy). Samples were collected either in a field expedition (labeled “Field”) or at the Cowichan Lake Research Station common garden in British Columbia, Canada (labeled “Cowichan”). The GPS coordinates listed for “Cowichan” samples are the collection origins of the seeds grown in the common garden, from which foliage samples were collected for this study.

| Species | Population | Sampling Label | Collection | Latitude | Longitude |
|---|---|---|---|---|---|
| *T. mertensiana* | 1 | TY | Field | 61° 3'19.73"N | 151°25'24.52"W |
| *T. mertensiana* | 2 | KEN | Field | 60°47'31.56"N | 149°12'43.70"W |
| *T. mertensiana* | 3 | Jun | Field | 58°16'50.35"N | 134°24'52.37"W |
| *T. mertensiana* | 4 | TER | Field | 54°55'30.00"N | 128°15'0.00"W |
| *T. mertensiana* | 5 | MD-MH | Field | 51°58'13.03"N | 118°28'23.71"W |
| *T. mertensiana* | 6 | SP-MH | Field | 51° 9'44.62"N | 118° 9'32.26"W |
| *T. mertensiana* | 7 | GP-MH | Field | 51°14'57.82"N | 117°28'50.68"W |
| *T. mertensiana* | 8 | AW-MH | Field | 49°14'17.29"N | 124°36'25.13"W |
| *T. mertensiana* | 9 | MS-MH | Field | 49°22'28.80"N | 122°57'24.16"W |
| *T. mertensiana* | 10 | A-MH-WA-02 | Field | 48°31'8.58"N | 120°40'27.19"W |
| *T. mertensiana* | 11 | A-MH-WA-01 | Field | 47°29'0.06"N | 123°37'1.45"W |
| *T. mertensiana* | 12 | MH-WA-02 | Field | 47°27'19.80"N | 121°26'24.00"W |
| *T. mertensiana* | 13 | MH-WA-01 | Field | 46°39'20.52"N | 121°24'30.96"W |
| *T. mertensiana* | 14 | MH-OR-05 | Field | 45°19'48.36"N | 121°42'28.44"W |
| *T. mertensiana* | 15 | A-MH-ID-01 | Field | 47°48'34.42"N | 116° 1'26.18"W |
| *T. mertensiana* | 16 | A-MH-MT-01 | Field | 47°25'50.02"N | 113°48'41.36"W |
| *T. mertensiana* | 17 | A-MH-ID-02 | Field | 46°28'51.89"N | 115°38'2.15"W |
| *T. mertensiana* | 18 | A-MH-OR-01 | Field | 45°21'4.25"N | 117°41'53.59"W |
| *T. mertensiana* | 19 | OLA | Field | 44° 7'13.99"N | 122° 5'46.99"W |
| *T. mertensiana* | 20 | MH-OR-02 | Field | 42° 5'20.76"N | 123°22'8.40"W |
| *T. mertensiana* | 21 | MH-CA-01 | Field | 41° 2'45.60"N | 122°47'52.44"W |
| *T. plicata* | 1 | N/A | Cowichan | 52°45'0.00"N | 131°50'60.00"W |
| *T. plicata* | 2 | N/A | Cowichan | 51°52'59.88"N | 127°46'0.12"W |
| *T. plicata* | 3 | N/A | Cowichan | 49°31'59.88"N | 124°57'0.00"W |
| *T. plicata* | 4 | N/A | Cowichan | 50°27'15.12"N | 122°15'0.72"W |
| *T. plicata* | 5 | N/A | Cowichan | 53°18'46.80"N | 120°15'21.60"W |
| *T. plicata* | 6 | N/A | Cowichan | 52°17'6.00"N | 119°10'47.28"W |
| *T. plicata* | 7 | N/A | Cowichan | 50°33'20.52"N | 117°27'29.88"W |
| *T. plicata* | 8 | N/A | Cowichan | 50°22'17.40"N | 119° 1'5.52"W |
| *T. plicata* | 9 | SC-WR | Field | 49°21'50.72"N | 120° 9'59.76"W |
| *T. plicata* | 10 | A-WC-WA-02 | Field | 48°30'9.90"N | 118°14'3.19"W |
| *T. plicata* | 11 | WC-WA-02 | Field | 47°33'5.76"N | 120°46'3.00"W |
| *T. plicata* | 12 | WC-WA-04 | Field | 47°31'9.48"N | 123°19'56.28"W |
| *T. plicata* | 13 | A-WC-ID-01 | Field | 47°54'20.02"N | 116°26'13.56"W |
| *T. plicata* | 14 | A-WC-MT-01 | Field | 48°10'3.47"N | 114° 0'11.95"W |
| *T. plicata* | 15 | A-WC-ID-04 | Field | 46°51'31.21"N | 115°38'45.20"W |
| *T. plicata* | 16 | A-WC-ID-03 | Field | 46°12'51.91"N | 115°32'12.88"W |
| *T. plicata* | 17 | SEL | Field | 46° 8'16.97"N | 114°24'50.72"W |
| *T. plicata* | 18 | N/A | Cowichan | 45°40'59.88"N | 123°25'0.12"W |
| *T. plicata* | 19 | WC-OR-03 | Field | 45°16'44.04"N | 121°43'55.20"W |
| *T. plicata* | 20 | N/A | Cowichan | 43°55'0.12"N | 122°46'0.12"W |
| *T. plicata* | 21 | WC-CA-01 | Field | 40°53'31.20"N | 123°56'32.64"W |
| *T. plicata* | 22 | WC-CA-02 | Field | 40°30'54.72"N | 124°16'28.20"W |

**Table S2.** Analysis of Molecular Variance (AMOVA)

Analysis of genetic variance among interior hemlock sampling sites and latitudinal-equivalent coastal sites (n = 14): (**A**) Variance between the north and south hemlock genetic lineages. (**B**) Variance between the coastal and interior distributions of hemlock. Analysis of genetic variance among interior redcedar sites and "Central" lineage coastal sites (n=17). (**C**) Variance between the "Central" and Idaho redcedar genetic lineages. (**D**) Variance between the coastal and interior distributions of hemlock, within the "Central" and Idaho genetic lineages. Asterisks indicate significant effects ($P < 0.05$).

| **A. North *vs.* South Hemlock Lineage** | | | | |
|---|---|---|---|---|
| Source of Variation | d.f. | Sum of Squares | Variance Components | Percent of Variation |
| Between Lineages | 1 | 108.71 | 1.11 | 5.81* |
| Among Sites Within Lineages | 12 | 329.93 | 0.93 | 4.87* |
| Among Individuals Within Sites | 87 | 2397.19 | 17.11 | 89.3* |
| **B. Coast *vs.* Interior Hemlock Mesic Region** | | | | |
| Source of Variation | d.f. | Sum of Squares | Variance Components | Percent of Variation |
| Between Regions | 1 | 39.18 | 0.04 | 0.22 |
| Among Sites Within Regions | 12 | 399.46 | 1.48 | 7.96* |
| Among Individuals Within Sites | 87 | 2397.19 | 17.11 | 91.83* |
| **C. Central *vs.* Idaho Redcedar Lineage** | | | | |
| Source of Variation | d.f. | Sum of Squares | Variance Components | Percent of Variation |
| Between Lineages | 1 | 14,338.16 | 107.35 | 3.52* |
| Among Sites Within Lineages | 15 | 77,154.64 | 16.68 | 0.55* |
| Among Individuals Within Sites | 102 | 500,831.00 | 1982.27 | 64.95* |
| Among Individuals | 119 | 112,522.50 | 945.56 | 30.98* |
| **D. Coast *vs.* Interior Redcedar Mesic Region** | | | | |
| Source of Variation | d.f. | Sum of Squares | Variance Components | Percent of Variation |
| Between Regions | 1 | 8,340.54 | 24.26 | 0.81* |
| Among Sites Within Regions | 15 | 83,152.27 | 45.24 | 1.51* |
| Among Individuals Within Sites | 102 | 500,831.00 | 1,982.27 | 66.13* |
| Among Individuals | 119 | 112,522.50 | 945.57 | 31.55* |

**Table S3.** Final *DIYABC* estimates for *Tsuga mertensiana* and *Thuja plicata* divergence times in number of generations (See Fig. 3). We assumed one generation to be equivalent to twenty years in both species for the purposes of this study.

| A. Mountain Hemlock | | | | | |
|---|---|---|---|---|---|
| Divergence | Mean | Median | Mode | 5th Percentile | 95th Percentile |
| t1 | 535 | 550 | 682 | 336 | 678 |
| t2 | 1,270 | 1,230 | 1,150 | 562 | 2,110 |
| t3 | 5,030 | 4,940 | 4,690 | 3,950 | 6,410 |
| **B. Western Redcedar** | | | | | |
| Divergence | Mean | Median | Mode | 5th Percentile | 95th Percentile |
| t1 | 1,180 | 1,170 | 1,090 | 509 | 1,910 |
| t2 | 3,010 | 2,910 | 2,650 | 2,110 | 4,260 |
| t3 | 3,530 | 3,330 | 2,890 | 2,390 | 5,540 |
| t4 | 7,000 | 6,960 | 7,040 | 4,880 | 9,240 |

**Table S4.** Published mountain hemlock pollen records.
Table of published data used to assemble Figure S2. Paleorecords listing mountain hemlock in the study region were classified into two categories based on the basal age of the sediment core and the first appearance of mountain hemlock pollen above a 1% threshold. Categorization was based on the paleorecord's published age model.

| Site Name | Latitude | Longitude | Publication | Pollen Age | Basal Age |
|---|---|---|---|---|---|
| Mineral Lake | 46.7262 | -122.1739 | Tsukada 1981 (8) | >20,000 | >20,000 |
| Kalaloch (approx.) | 47.634 | -124.3825 | Heusser, 1972 (9) | >20,000 | >20,000 |
| Little Lake | 44.17 | -123.58 | Worona & Whitlock, 1995 (10) | >20,000 | >20,000 |
| Grass Lake | 41.65 | -122.17 | Hakala & Adam, 2004 (11) | >20,000 | >20,000 |
| Fargher Pond | 45.89 | -122.52 | Grigg & Whitlock, 2002 (12) | >20,000 | >20,000 |
| Humptulips Bog | 47.17 | -123.55 | Heusser et al, 1999 (13) | >20,000 | >20,000 |
| Battleground Lake | 45.8 | -122.49 | Whitlock, 1992; Barnosky, 1985 (14, 15) | >20,000 | >20,000 |
| Moose Lake | 47.88 | -123.35 | Gavin et al, 2001 (16) | >20,000 | >20,000 |
| Lake Washington | 47.6733 | -122.2233 | Leopold et al. 1982 (17) | >15,000 | >15,000 |
| Top Lake | 54.48 | -130.95 | McLaren, 2008 (18) | >15,000 | >15,000 |
| Mosquito Lake Bog | 48.77 | -122.12 | Steventon & Kutzbach, 1986; Hansen & Easterbrook, 1974 (19, 20) | >15,000 | >15,000 |
| Misty Lake | 50.6 | -127.25 | Lacourse, 2005 (21) | >15,000 | >15,000 |
| Indian Prairie Fen | 44.63 | -122.58 | Sea & Whitlock, 1995 (22) | >15,000 | >15,000 |
| Little Willow Lake | 40.41 | -121.39 | West, 2003 (23) | >15,000 | >15,000 |
| Pixie Lake | 48.6 | -124.2 | Brown & Hebda, 2002; Brown & Hebda, 2003 (24, 25) | >15,000 | >15,000 |
| Pleasant Island | 58.21 | -135.37 | Hansen & Engstrom, 1996 (26) | >10,000 | >15,000 |
| Bear Cove Bog | 50.716 | -127.46 | Hebda 1983 (27) | >10,000 | >15,000 |
| Porphyry Lake | 48.91 | -123.83 | Brown & Hebda, 2002; Brown & Hebda, 2003 (24, 25) | >10,000 | >15,000 |
| Marion Lake | 49.308 | -122.547 | Mathewes 1973 (28) | >10,000 | >10,000 |
| Tiny Lake | 51.11 | -127.22 | Galloway et al., 2009 (29) | >10,000 | >10,000 |
| Yahoo Lake | 47.68 | -124.02 | Gavin et al, 2013 (30) | >10,000 | >10,000 |
| Martins Lake | 47.71 | -123.53 | Gavin et al, 2001 (16) | >10,000 | >10,000 |
| Kirk Lake | 48.23 | -121.62 | Cwynar, 1987 (31) | >10,000 | >10,000 |
| Killebrew Lake Fen | 48.61 | -122.9 | Leopold et al, 2016 (32) | >10,000 | >10,000 |
| Mono Lake | 38.01 | -119.03 | Davis, 1999 (33) | >10,000 | >10,000 |
| Cedar Lake | 41.2 | -122.5 | West, 1989 (34) | >10,000 | >10,000 |
| Gold Lake Bog | 43.65 | -122.04 | Sea & Whitlock, 1995 (22) | >10,000 | >10,000 |
| Tumalo Lake | 44.02 | -121.54 | Long et al, 2011 (35) | >10,000 | >10,000 |

| | | | | | |
|---|---|---|---|---|---|
| Fishblue (Blue) Lake | 49.99 | -121.49 | Mathewes & Westgate, 1980; Mathewes & King, 1989 (36, 37) | >10,000 | >10,000 |
| Cassiope Pond | 50.17 | -127.75 | Hebda & Allen, 1997 (38) | >10,000 | >10,000 |
| Pyrola Lake | 50.18 | -127.7 | Hebda, 1997 (39) | >10,000 | >10,000 |
| Woods Lake | 51 | -127.27 | Stolze et al, 2007; Stolz, 2004 (40, 41) | >10,000 | >10,000 |
| East Sooke Fen | 48.35 | -123.68 | Brown & Hebda, 2002; Brown & Hebda, 2003 (24, 25) | >10,000 | >10,000 |
| Walker Lake | 48.53 | -124 | Brown & Hebda, 2002; Brown & Hebda, 2003 (24, 25) | >10,000 | >10,000 |
| Surprise Lake | 49.32 | -122.56 | Mathewes, 1973 (28) | >10,000 | >10,000 |
| Two Frog Lake | 51.11 | -127.53 | Galloway et al, 2007 (42) | >10,000 | >10,000 |
| Louise Pond | 52.95 | -131.76 | Pellatt & Mathewes, 1994; Pellatt & Mathewes, 1997 (43, 44) | >10,000 | >10,000 |
| Mitkof Island | 56.81 | -132.94 | Ager et al, 2010 (45) | >10,000 | >10,000 |
| Pine Lakes Muskeg | 59.54 | -139.57 | Peteet 1991 (46) | >5,000 | >10,000 |
| Drizzle Pond | 59.7 | -135.09 | Spear and Cwynar 1997 (47) | >5,000 | >10,000 |
| Lily Lake | 59.2 | -135.4 | Cwynar, 1990 (48) | >5,000 | >10,000 |
| Pinecrest Lake | 49.49 | -121.43 | Mathewes & Rouse, 1975; Mathewes et al, 1973 (28, 49) | >5,000 | >10,000 |
| Tioga Pass Pond | 37.91 | -119.26 | Anderson, 1987; Anderson, 1990 (50, 51) | >5,000 | >10,000 |
| Barrett Lake | 37.6 | -119.01 | Anderson, 1987; Anderson, 1990 (50, 51) | >5,000 | >10,000 |
| Berendon Fen | 56.24 | -130.06 | Clague & Mathewes, 1996; Clague et al, 2004 (52, 53) | >5,000 | >5,000 |
| Shangri-La Bog | 53.27 | -132.41 | Pellatt & Mathewes, 1997 (44) | >5,000 | >5,000 |
| Hidden Lake | 60.49 | -150.30 | Ager & Wright, 1983; Ager & Sims, 1982; Ager & Brubaker, 1985; Rymer & Sims, 1982 (54–57) | >0 | >15,000 |
| Waterdevil Lake | 59.76 | -134.93 | Spear and Cwynar 1997 (47) | >0 | >10,000 |
| Icy Cape | 59.57 | -141.24 | Peteet, 1986 (58) | >0 | >10,000 |
| Golden | 60.61 | -147.149 | Heusser, 1983 (59) | >0 | >10,000 |
| Munday Creek | 60.03 | -141.97 | Peteet, 1986 (58) | >0 | >10,000 |
| Bluff Lake | 41.35 | -122.56 | Mohr et al, 2000; Mohr, 1997 (60, 61) | >0 | >10,000 |
| Starkweather Pond | 37.66 | -119.07 | Anderson, 1987; Anderson, 1990 (50, 51) | >0 | >10,000 |
| Dismal Lake | 47.12 | -115.63 | Herring et al, 2018 (62) | >0 | >10,000 |
| Rocky Ridge Lake | 46.44 | -115.49 | Herring et al, 2018 (62) | >0 | >10,000 |
| Horseshoe Lake | 46.55 | -115.07 | Herring et al, 2018 (62) | >0 | >10,000 |
| Testalinden Lake | 49.12 | -119.68 | Heinrichs et al, 2001 (63) | >0 | >5,000 |
| Eagle Lake | 51.04 | -118.15 | Rosenberg et al, 2003 (64) | >0 | >5,000 |
| Spillway Pond | 56.24 | -130.07 | Clague & Mathewes, 1996; Clague et al, 2004 (52, 53) | >0 | >0 |
| Moraine Bog | 51.33 | -124.9 | Arsenault et al, 2007 (65) | >0 | >0 |

1. S. T. Kalinowski, How well do evolutionary trees describe genetic relationships among populations? *Heredity* **102**, 506–513 (2009).

2. B. S. Weir, C. C. Cockerham, Estimating F-statistics for the analysis of population structure. *evolution* **38**, 1358–1370 (1984).

3. J.-M. Cornuet, *et al.*, DIYABC v2. 0: a software to make approximate Bayesian computation inferences about population history using single nucleotide polymorphism, DNA sequence and microsatellite data. *Bioinformatics* **30**, 1187–1189 (2014).

4. D. G. W. Edwards, C. L. Leadem, The reproductive biology of western red cedar with some observations on nursery production and prospects for seed orchards (1988).

5. D. Minore, Thuja plicata Donn ex D. Don—western redcedar. *Silvics of North America* **1**, 590–600 (1990).

6. D. P. Turner, Successional relationships and a comparison of biological characteristics among six northwestern conifers. *Bulletin of the Torrey Botanical Club*, 421–428 (1985).

7. J. E. Means, Tsuga mertensiana (Bong.) Carr. mountain hemlock. *Silvics of North America* **1**, 623–631 (1990).

8. M. Tsukada, S. Sugita, D. M. Hibbert, Paleoecology in the Pacific Northwest. I. Late Quaternary vegetation and climate. *Proceedings-International Association of Theoretical and Applied Limnology* (1981).

18. D. McLaren, "Sea level change and archaeological site locations on the Dundas Island Archipelago of North Coastal British Columbia." (2008).

40. S. Stolze, "A record of late Quaternary vegetation and climate change from Woods Lake, Seymour Inlet, coastal British Columbia, Canada," Carleton University. (2004).

50. R. S. Anderson, Late-Quaternary Environments of the Sierra Nevada, California. Ph.D. Thesis, University of Arizona. (1987).

61. J. A. Mohr, "Postglacial vegetation and fire history near Bluff Lake, Klamath Mountains, California," University of Oregon. (1997).

62. E. M. Herring, D. G. Gavin, S. Z. Dobrowski, M. Fernandez, F. S. Hu, Ecological history of a long-lived conifer in a disjunct population. *Journal of Ecology* (2017).

63. M. L. Heinrichs, R. J. Hebda, I. R. Walker, Holocene vegetation and natural disturbance in the Engelmann Spruce Subalpine Fir biogeoclimatic zone at Mount Kobau, British Columbia. *Canadian journal of forest research* **31**, 2183–2199 (2001).

64. S. M. Rosenberg, I. R. Walker, R. W. Mathewes, Postglacial spread of hemlock (Tsuga) and vegetation history in Mount Revelstoke National Park, British Columbia, Canada. *Canadian Journal of Botany* **81**, 139–151 (2003).

65. T. A. Arsenault, J. J. Clague, R. W. Mathewes, Late Holocene vegetation and climate change at Moraine Bog, Tiedemann Glacier, southern Coast Mountains, British Columbia. *Canadian Journal of Earth Sciences* **44**, 707–719 (2007).
